# Tree phylogenetic diversity supports nature’s contributions to people, but is at risk from human population growth

**DOI:** 10.1101/2021.02.13.430985

**Authors:** T. Jonathan Davies, Olivier Maurin, Kowiyou Yessoufou, Barnabas H. Daru, Bezeng S. Bezeng, Hanno Schaefer, Wilfried Thuiller, Michelle van der Bank

## Abstract

There is growing evidence for a link between biodiversity and ecosystem function, and for a correlation between human population and the species diversity of plants and animals in a region. Here, we suggest these relationships might not be independent. Using a comprehensive phylogeny of southern African trees and structural equation modelling, we show that human population density correlates with tree phylogenetic diversity and show that this relationship is stronger than the correlation with species richness alone. Further, we demonstrate that areas high in phylogenetic diversity support a greater diversity of ecosystem goods and services, indicating that the evolutionary processes responsible for generating variation among living organisms are also key to the provisioning of nature’s contributions to people. Our results raise the intriguing possibility that the history of human settlement in southern Africa may have been shaped, in part, by the evolutionary history of its tree flora. However, the correlation between human population and tree diversity generates a conflict between people and nature. Our study suggests that future human population growth may threaten the contributions to people provided by intact and phylogenetically diverse ecosystems.

---

Human population has increased dramatically over the last 200 years, and is predicted to approach 9 billion by 2050 (United Nations 2004). The distribution of people, however, is highly uneven. In 2008, the urban population outnumbered the rural population for the first time (United Nations 2011), and in much of the developed world the human population is stable or decreasing (Murray *et al.* 2018). In contrast, although urbanizing rapidly, the human population in sub-Saharan Africa is still increasing (Murray *et al.* 2018), and remains predominantly rural (UNICEF 2012), reflecting a closer relationship between the natural environment and people. Much of the human population in sub-Saharan Africa depends directly or indirectly on the goods and services provided by biodiversity (Shackleton *et al.* 2008; Egoh *et al.* 2012), including clean water, soil productivity, firewood/charcoal, game animals, and medicinal plants (Daily & others 1997; Postel *et al.* 2012) – nature’s contributions to people (Díaz *et al.* 2018; Brondizio *et al.* 2019). However, many of these services are being degraded through habitat loss and transformation, exacerbated by rapid urbanization, agricultural expansion, ‘land grabbing’ by foreign nations for food and biofuel production, and climate change (Egoh *et al.* 2012; Malherbe *et al.* 2019). The poor, and rural communities, tend to be most impacted by losses of ecosystem services as they frequently lack alternatives (Hope Sr 2007; United Nation Economic Commission for Africa 2010; Egoh *et al.* 2012; Kumar & Yashiro 2014), while poverty is a major underlying contributor to environmental degradation (World Commission on Environment and Development 1987; Olanipekun *et al.* 2019), although the relationship is complex (Barbier & Hochard 2018).

Quantifying the value of ecosystem services is challenging because many such services are difficult to measure and different value systems do not necessarily align closely (Costanza *et al.* 2017; Díaz *et al.* 2018; Brondizio *et al.* 2019), although estimates suggest their crude economic worth might be large (Costanza *et al.* 1998). Species richness provides one surrogate measure of ecosystem services that is easily quantifiable. The link between species richness and ecosystem function has been the focus of much attention (Hooper *et al.* 2005; Cardinale *et al.* 2012; Hooper *et al.* 2012), and experimental manipulations suggest that, in some systems, high species richness might confer both greater ecosystem productivity and stability (Tilman *et al.* 2012; Isbell *et al.* 2015), and also support a greater diversity of services (Hector & Bagchi 2007, see also Gamfeldt *et al.*, 2013, for an ecosystem example). However, the correlation between ecosystem services and species richness at the landscape scale is often mixed (Egoh *et al.* 2009; O’Farrell *et al.* 2010; Manhães *et al.* 2016, see also Anderson *et al.* 2009). This discrepancy may be explained by the mechanisms underlying biodiversity-ecosystem function relationships. Species richness is thought to enhance ecosystem function and stability, at least in part, via species complementarity (Loreau & Hector 2001), which assumes that different species fulfil different functions and occupy separate ecological niches. However, simple measures of species richness may not accurately capture the ecological variation among species.

Phylogenetic diversity – the summed branch lengths that connect species on a phylogenetic tree – provides a more inclusive metric for quantifying species differences (Faith 1992). Because we expect species to diverge over time in their ecological and physiological traits, the set of species with higher phylogenetic diversity is predicted to capture greater feature diversity (Faith 1992, but see Tucker *et al.* 2018; Mazel *et al.* 2018). Experimental data have demonstrated a link between phylogenetic diversity and ecosystem properties (e.g. productivity and stability), which can be stronger than observed for species richness alone (Cadotte *et al.* 2008, but see Venail *et al.* 2015). There is also evidence that phylogenetic diversity contributes to forest productivity (Hao *et al.* 2018, see also Paquette & Messier 2011). Forest *et al.* (2007) additionally demonstrated that a greater phylogenetic diversity might translates into a greater diversity of ecosystem services, using data on medicinal plants within the Cape of South Africa biodiversity hotspot.

Here, we evaluate the relationship between tree phylogenetic diversity, provisioning of ecosystem services – nature’s contributions to people – and human population density in southern Africa. In Africa and globally, trees have supported and sustained human life through provisioning of various regulating (e.g. carbon sequestration, soil preservation and clean air), provisioning (e.g. food, firewood and charcoal and medicine) and cultural services (Daily & others 1997; Millennium Ecosystem Assessment 2005), worth many millions of dollars (see Commonwealth Science Council and Food and Agriculture Organization 1993). Some of these services provide global benefits (e.g. many regulating services), but others have more local benefits (especially provisioning and cultural services). If such services are linked to species functional traits (Cadotte *et al.* 2011; Huang *et al.* 2019), we would predict that the diversity of ecosystem services within a region would be a product of not only species number, but also of how different species are from one another – i.e. their phylogenetic diversity (Forest *et al.* 2007; Faith *et al.* 2010). We explore this hypothesis here, using a comprehensive phylogeny for the trees of southern Africa (Fig. 1) – a regional flora well known for its incredible plant diversity.

**Figure 1.**
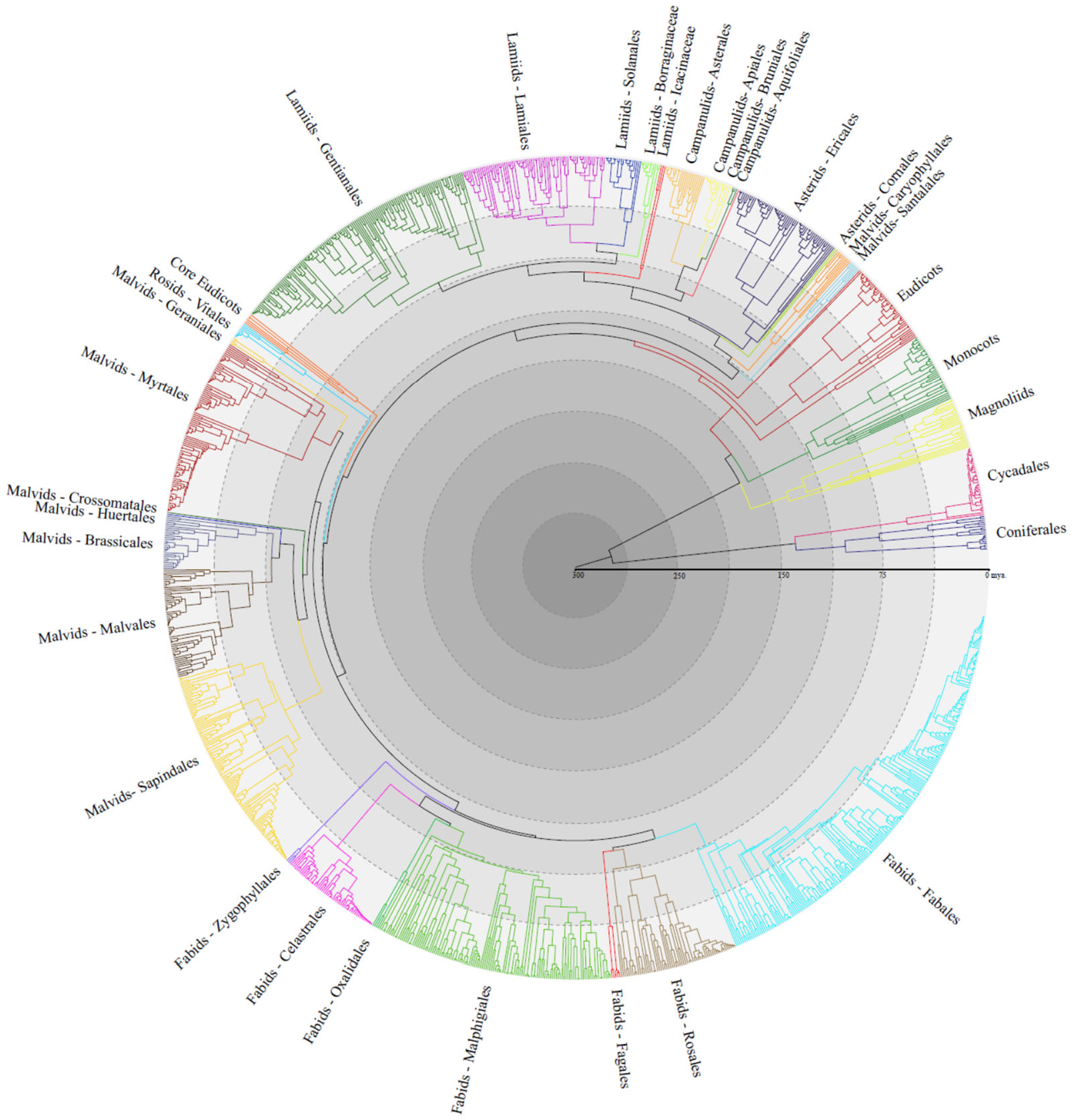
Phylogeny of 1,308 southern African tree species. Reconstruction based on DNA barcodes for land plants using maximum likelihood and transforming branch lengths to millions of years by enforcing a relaxed molecular clock and multiple fossil calibrations. Colours indicate higher-level taxonomic groupings.

## RESULTS AND DISCUSSION

We first evaluated the statistical linkages between human population density and the natural distribution of native woody plant (tree) richness and phylogenetic diversity across southern Africa. We used a phylogenetic tree encompassing 2,486 tree species in the region, spanning 115 families and 541 genera, and reconstructed using DNA barcode sequences for the core plant barcodes (*rbcL*a and *matK*) (CBOL Plant Working Group 2009) generated by the authors as part of a six year-long data collection effort (for additional details see Maurin *et al.* 2014 and Charles-Dominique *et al.* 2016). Hotspots of tree species diversity are distributed along the coastal regions of eastern South Africa through to south Mozambique and Zimbabwe (Fig. 2A, B), and differ notably from richness maps of total plant diversity for the region (> 20,000 species), which emphasize the high species richness of the Cape Floristic Kingdom (Cowling & Hilton-Taylor 1994) with its extraordinary radiations in a few (non-tree) clades (e.g. *Erica*, Restionaceae and Iridaceae).

**Figure 2.**
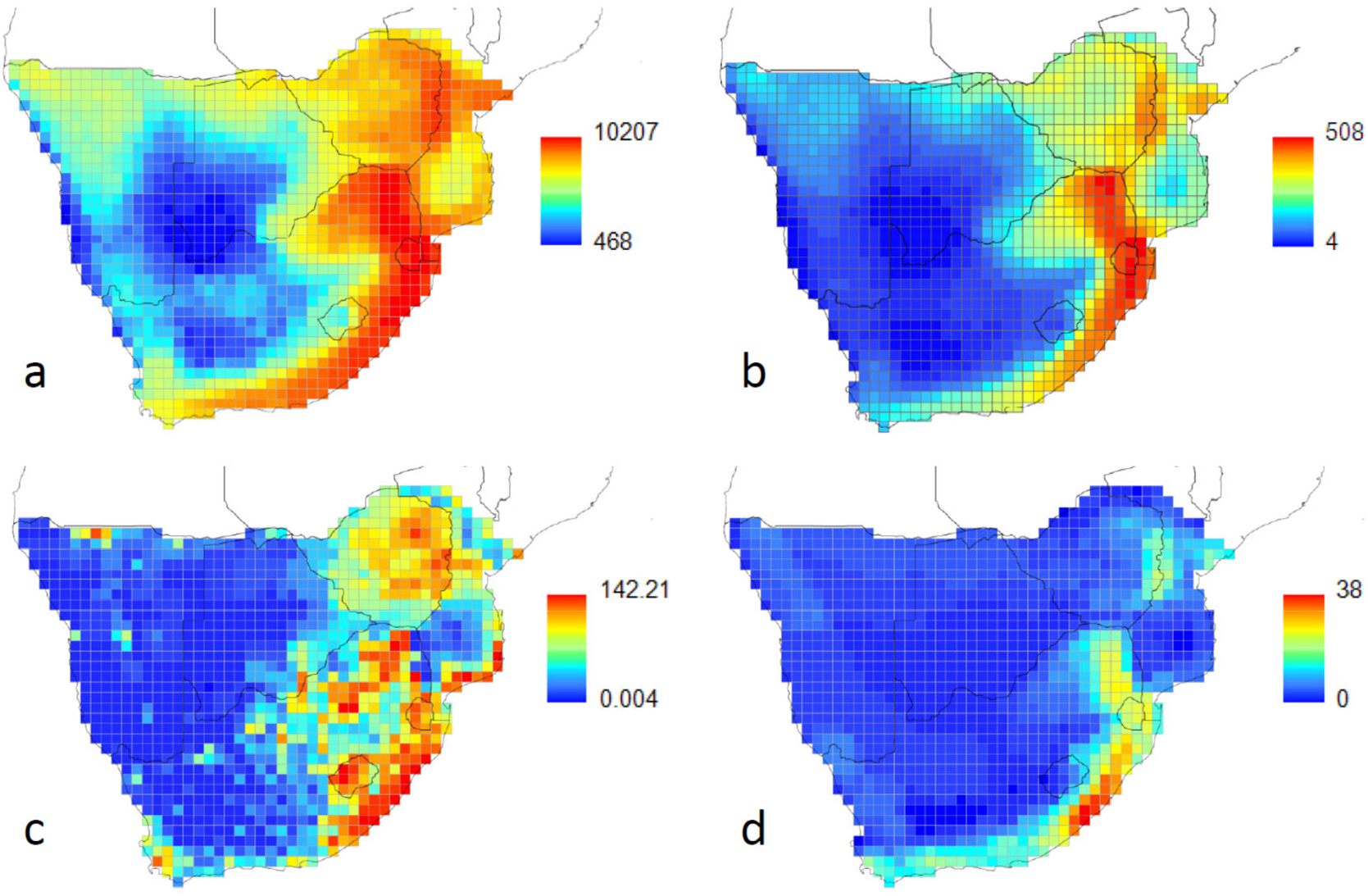
Geographical distribution of tree diversity and human population density. Tree phylogenetic diversity (myr) (**a**), total tree species richness (**b**), mean human population density (persons/km^2^) excluding urban areas (**c**), and richness of threatened trees (**d**). Data were divided into 32 quantiles and plotted on a 0.5° × 0.5° grid.

### A covariation between people and tree phylogenetic diversity

We reveal a significant and strong positive correlation between human population density and tree phylogenetic diversity (Pearson’s r = 0.76, p<0.01 on 7.73 adjusted degrees of freedom to correct for spatial non-independence, Dutilleul *et al.* 1993; Figs. 2A - D), and show that this relationship is stronger than the correlation with species richness alone (r = 0.62 and p<0.01 on 16.74 adjusted degrees of freedom). A positive covariation between wildlife species richness and human population has been noted previously (Balmford *et al.* 2001; Chown *et al.* 2003; Evans *et al.* 2006; Luck 2007) but, to the best of our knowledge, we are the first to show that the correlation with phylogenetic diversity is stronger. Our results thus indicate that human population density co-varies not only with the number of tree species but also with how different they are from each other, as reflected in their evolutionary history.

One explanation for the association between human population density and the phylogenetic diversity of trees is that both covary with correlated environmental variables (Chown *et al.* 2003; Evans *et al.* 2006; Luck 2007). For example, tree diversity might correlate with environmental productivity and regions suitable for agriculture and subsistence. We therefore used Structural Equation Models (SEMs) to describe the direct and indirect effect of environmental energy (here, modeled as actual evapotranspiration, AET), topographic heterogeneity (indexed by elevation), and tree species diversity (i.e. richness and phylogenetic diversity) on human population density. While topographic heterogeneity was not a significant predictor in our models, environmental energy had a direct effect on both human population density and PD. However, the effect of AET on human population density was mediated by an indirect effect via phylogenetic diversity, and the direct effect of tree phylogenetic diversity on human density (β=0.43) was greater than the direct effect of AET (β=0.34) (Fig. 3A).

**Figure 3.**
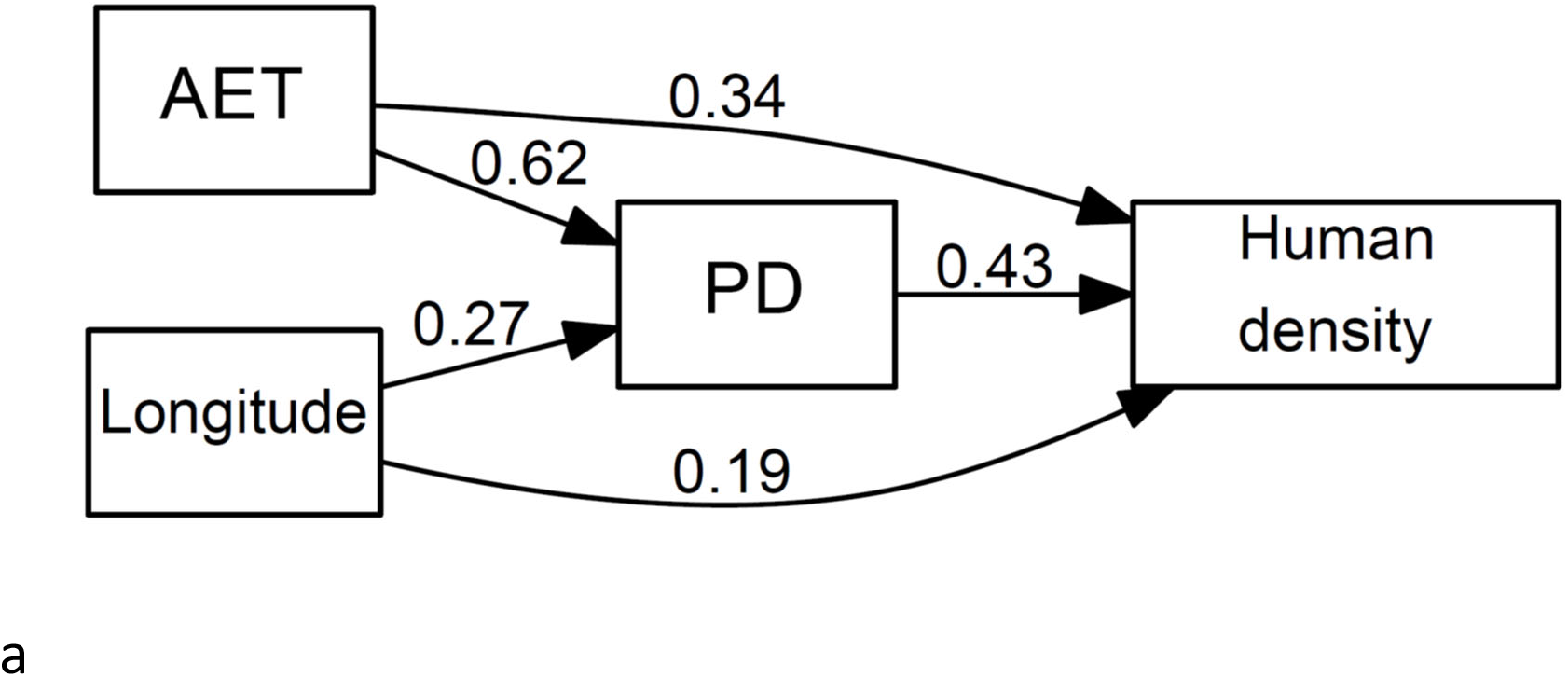
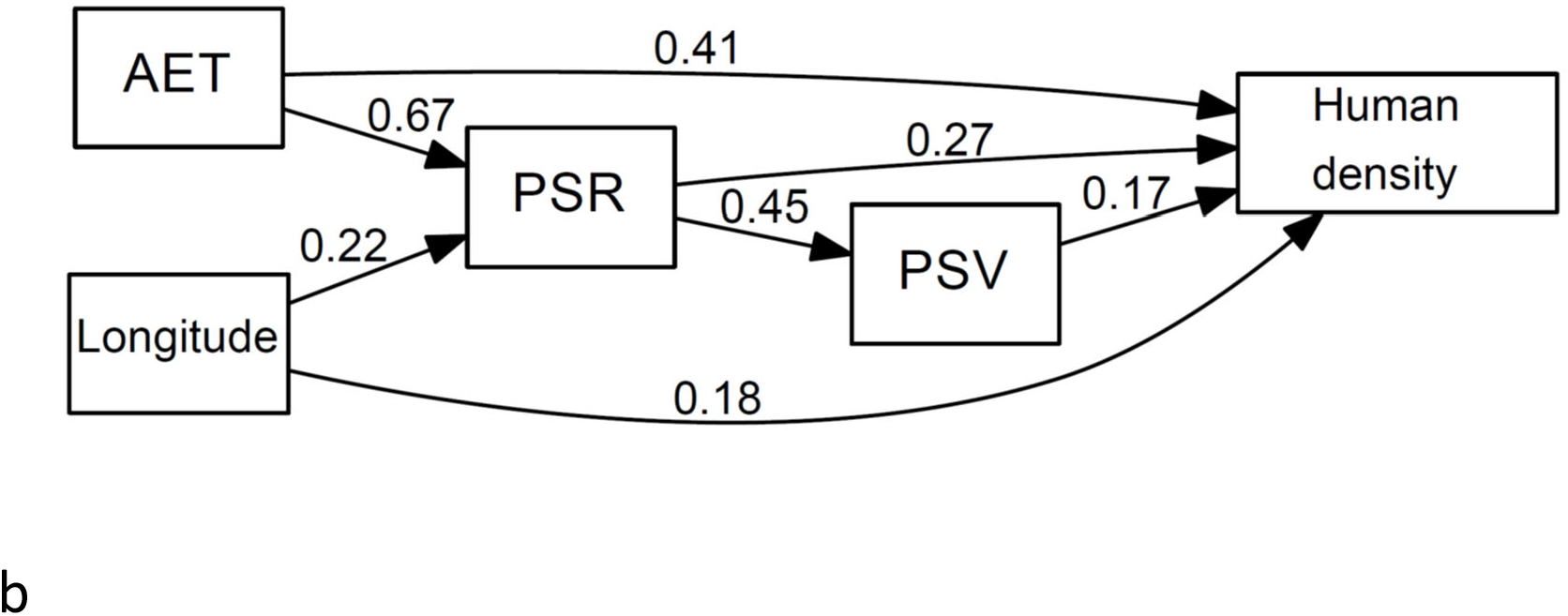
Structural Equation Models. Illustration of the direct and indirect effects of actual evapotranspiration (AET), phylogenetic diversity (PD) and longitude on human population density (**a**), and an equivalent model decomposing phylogenetic diversity into a richness independent measure of evolutionary disparity (PSV) and a measure of richness discounted for species relatedness (PSR) (**b**). Model A is favored by model fit statistics (BIC = 1129.78 and 1275.75 for model A and B respectively), although model B explains more of the total variation in human population density (r^2^ = 0.56 and 0.59 for human density, model A and B respectively). Arrows indicate inferred direction of causation and numbers relative path strengths (standardized coefficients). The inclusion of additional pathways, such as the direct and indirect effects of elevation and latitude (Fig. S4), was not supported by model comparison statistics.

To separate out the importance of number of species versus species evolutionary disparity, we calculated a measure of evolutionary dispersion independent of richness, PSV, and a separate measure of richness discounted for species phylogenetic relatedness, PSR (see Helmus *et al.* 2007). The direct effect of PSR (the richness component) on human density was weaker than observed for total phylogenetic diversity, with the missing variation explained by PSV (the phylogenetic component) (Fig. 3B). However, we note that the SEM including phylogenetic diversity is favored overall (BIC = 1129.78 and 1275.75 for the SEMs with phylogenetic diversity and PSR plus PSV, respectively), as it captures both richness and evolutionary disparity in a single unifying metric. The independent contribution of phylogenetic diversity is further demonstrated when we include in our SEM both species richness and the residual variation in phylogenetic diversity from the regression of phylogenetic diversity against richness (Fig. S1).

There are, of course, many additional predictor variables that we did not explore in our models. For example, in southern Africa, edaphic factors might be particularly important for plant diversity (Goldblatt 1997; Goldblatt & Manning 2002). However, high regional plant species diversity is often associated with nutrient poor soils, such as in the fynbos biome (Cowling *et al.* 1996), presumably poor for agriculture, and which may have limited early human settlement of the Cape peninsular (Cowling *et al.* 1996). In addition, it is not obvious how environmental or climatic factors could drive the correlation between human population and tree phylogenetic diversity independent of tree species richness.

### Tree phylogenetic diversity supports nature’s contributions to people

We evaluated the relationship between tree phylogenetic diversity and the provisioning of ecosystem goods and services – nature’s contributions to people. We examined the diversity of ecosystem services using data on 11 classes of recorded uses, including cultural, medicinal, forage, and food. Because limited availability of data on species’ uses precluded the use of traditional metrics for capturing functional diversity (e.g. Petchey & Gaston 2002 and Laliberté & Legendre 2010), we used a simulation approach. Our method quantified the median number of species required to represent at least once each of the different uses identified for the species within a given 0.5° grid cell when drawing species at random from the species set. Overall, more species had to be sampled to represent the same number of uses in more species-rich cells, suggesting functional redundancy across species; however, after correcting for variation in species richness, new uses were encountered at an increased rate in more phylogenetically dispersed communities (Table 1). More phylogenetically diverse tree communities therefore tend to represent a greater number of ecosystem services than tree communities of equal richness but lower phylogenetic diversity.

**Table 1.** Regression of median number of species required to capture the uses represented within each grid cell.

| Model | $r^2_{\text{adjusted}}$ | DF | P-value | Coefficient | Estimate<br>(s.e.) | t-value |
| --- | --- | --- | --- | --- | --- | --- |
| 1 | 0.77 | 1204 | <0.001 | Log(PSR)* | 4.97 (0.97) | 59.43 |
|  |  |  |  | PSV* | -29.71 (2.73) | -10.90 |
| 2 | 0.78 | 1204 | <0.001 | Log(Richness)* | 4.75 (0.07) | 65.16 |
|  |  |  |  | PD<sub>residuals</sub>* | -0.002<br>(<0.01) | -11.59 |

\*significant at P<0.001

It is tempting to speculate on the causal linkages between the provisioning of nature’s contributions to people, tree phylogenetic diversity and human population. For example, it is possible that the correlations we reveal reflect in some part the settlement of people in Africa historically, which may have been influenced by the ecosystem services that more phylogenetically diverse tree communities provided, perhaps as part of a shift toward more settled agriculture. However, it is also likely that over the history of human settlement in Africa, humans have shaped the local environment to their needs, including the planting of some species outside their natural range. Our range maps were for native distributions, and at the regional scale of our analysis it thus seems improbable that humans have shaped floristic biodiversity gradients across sub-Saharan Africa. Nonetheless, we cannot exclude this possibility, and the SEM for either direction of causality is equally as well-supported (BIC = 1129.78 for both modeled pathways).

### A conflict between trees and people

Irrespective of the underlying drivers, the correlation between human population density and tree diversity presents a potential conflict between people and nature (Balmford *et al.* 2001), especially, as demand for energy and land for agriculture intensifies. Using information on the extinction risk status from the Red List for South African plants (http://redlist.sanbi.org/), we show that not only does the total number of tree species correlate positively with human population, but so too does the number of threatened species (r = 0.54, p<0.001). It is likely that as human population in Africa becomes more urbanized, and as species diversity decreases through local and global extinctions, the relationship between human population density and species diversity will be eroded (Fjeldså & Burgess 2008).

There has been increasing emphasis on the preservation of phylogenetic diversity in the conservation literature (Purvis *et al.* 2005; Forest *et al.* 2007; Pollock *et al.* 2017; Davis *et al.* 2018). Our study suggests how the failure to protect phylogenetic diversity might reduce the provisioning of natural goods and services, likely with disproportion impact on the rural poor – those least able to afford alternatives.

## METHODS

Over a six year-long data collection effort, we sampled 1,306 tree species (defined as woody plants with stems or pseudostems > 0.5 m in height) representing 115 families and 541 genera, across southern Africa (Appendix S1). This region encompasses an area of over 4 million km^2^ located between 15°38’ S and 34°50’ S, and 11°45’ E and 36°29’ E, and includes Botswana, Lesotho, Mozambique (south of the Zambezi river), Namibia, South Africa, Swaziland, and Zimbabwe. We sequenced the core plant barcodes (*rbcL*a and *matK*) (CBOL Plant Working Group 2009) from sampled material, and merged these data with matching barcode sequence data in GenBank from tree species native to the region. Voucher specimen information and GenBank accession numbers are listed in Appendix S2 and on the BOLD DataSystem (www.boldsystems.org).

Phylogenetic reconstruction followed the protocol detailed in Maurin *et al.* (2014). Branch lengths were calibrated in millions of years using Bayesian MCMC implemented in BEAST v.1.4.8 (Drummond & Rambaut 2007) and 28 fossil calibrations from Bell *et al.* (2010) (Table S1) as minimum age constrains, except for the root of the Eudicots, which was set at 124 myr.

Species distribution data were extracted from Coates Palgrave *et al.* (2002) and Van Wyk *et al.* (2011), and overlaid on a 0.5° × 0.5° grid (approximately 50 km × 50 km). For each species, we then collated data on 11 separate uses: firewood and charcoal, carving, building and structural, spiritual, cultural, food, ornamental, forage and fodder, shade, chemical compounds, and medicinal, plus threat category from the IUCN Red Data List of Threatened Species (http://redlist.sanbi.org/). All data and source references are listed in Appendix S3. Geographical information system (GIS) data were obtained for actual evapotranspiration, AET (dataset GNV183; http://www.grid.unep.ch/data/), as a proxy for productivity, elevation (dataset GTOPO30; http://www1.gsi.go.jp/geowww/globalmap-gsi/gtopo30/gtopo30.html), and human population density (dataset GPWv3 http://sedac.ciesin.columbia.edu/data/collection/gpw-v3). Data processing was performed in ArcMap v.10.0 and R (v. 3.5.0) using the libraries *ape, picante, sf, raster, rgdal, maptools, maps, sp*, and *ggplot2*. Because human population density can be dramatically elevated in urban centres, we excluded outliers, classified as cells with z-scores >3, prior to estimating average population densities. Our population outliers identified cells with human population density >489 people/km^2^, matching closely accepted criteria for urban areas in South Africa (≥500 people/km^2^) (Statistics South Africa 2001).

To evaluate the relationships between human population and tree diversity, we first correlated tree species richness and phylogenetic diversity (PD – sum of the phylogenetic branch lengths excluding the root branch; Faith 1992) against human population density, adjusting degrees of freedom to correct for spatial non-independence (Dutilleul *et al.* 1993). We report results using half-degree cells because this represents the native coordinate system of the climate and human population density layers; however, correlations using equal-area grid cells were highly similar (slope = 0.39±0.01 and 0.36±0.01, t = 42.75 and 45.22 for the regression of log tree species richness against human population density across 0.5° × 0.5° grid cells and on an equal area 50 km × 50 km grid, respectively).

We excluded urban areas with high human population density from our analysis (see above) because other factors, such as economic development, industrialisation and global trade have likely influenced the geography of urban population densities. However, we note that cities in Africa predate European colonialization (Mboup 2019), and the correlations with phylogenetic diversity are largely unchanged when centres of high population density were included; Fig. S2).

We used structural equation modelling (SEM) to describe the direct and indirect effects of environment and tree diversity on human population. Before model fitting, we first quantified the pairwise correlations strengths between human population density, AET, tree species richness and PD, variance in elevation, latitude and longitude, respectively (Fig. S3). We then used the *sem* r-library (Fox 2006) to model structural relationships, including only pathways with pairwise correlation strengths >0.5 to avoid issues of colinearity. To account for spatial structure in the model, we additionally considered the direct effects of longitude and latitude. All variables except latitude and longitude were log-transformed prior to analysis. The inclusion of additional pathways, such as the direct and indirect effects of elevation and latitude (Fig. S4), were not supported by model comparison statistics (BIC = 1411.17 versus 1129.78 for the more complex model and the reduced model described above). To separate out the importance of number of species versus species evolutionary disparity, we additionally jointly modelled a richness independent component of evolutionary disparity, PSV, equivalent to the mean pairwise distance between taxa in the assemblage, and a separate measure of richness discounted for species relatedness, PSR. Last, to evaluate model sensitivity, we constructed matching SEMs but included species richness and the residual variation in PD from the regression of PD against richness in place of PSR and PSV, respectively.

We used a simulation approach to test whether more phylogenetically diverse sets of species tended to capture a greater diversity of uses (ecosystem services/nature’s contributions to people). Because we had comprehensive data on only a limited number of uses (Appendix S3), it was not possible to simply correlate diversity of services against diversity of species (all uses were represented by at least one tree in the majority of cells). Therefore, for each cell, we randomly re-sampled species without replacement, and recorded the median number of species required to represent at least once all uses listed for the entire species set in the cell (1000 randomizations). We then regressed the median richness required for capturing diversity of recorded uses against per-cell PSV plus PSR, and in a second model per-cell species richness plus residual PD.

## Acknowledgements

We thank the Government of Canada through Genome Canada and the Ontario Genomics Institute (2008-OGI-ICI-03), the University of Johannesburg, the South African National Research Foundation, the Royal Society (U.K.), Toyota South Africa through the Toyota Enviro Outreach Programme for financial support and various local and international authorities granting us plant collections permits. KY is supported by NRF grant #112113.

## Author Contributions

M.vd.B., O.M. and T.J.D. conceived and designed the experiments. O.M., B.H.D., B.S.B. and M.vd.B. performed the experiments. T.J.D., O.M. and H.S. analyzed the data. M.vd.B., H.S. and W.T. contributed reagents/materials/analysis tools. T.J.D., K.Y., B.H.D., W.T., O.M. and M.vd.B. wrote the paper.

